# Accuracy and completeness of long read metagenomic assemblies

**DOI:** 10.1101/2022.11.23.517703

**Authors:** Jeremy Buttler, Devin Drown

## Abstract

Microbes, we can learn how microbes influence the surrounding environment, contribute to human health, and understand which pathogen interactions result in differences in disease severity. Metagenomics can be used as a tool to explore the interactions between microbes. Metagenomic assemblies built using long read nanopore data depend on the read level accuracy. The read level accuracy of nanopore sequencing has made dramatic improvements over the past several years. However, we do not know if the increased read level accuracy allows for faster assemblers to make as accurate metagenomic assemblies as slower assemblers. Here, we present the results of a benchmarking study comparing three commonly used long read assemblers, Flye, Raven, and Redbean. We used a prepared DNA standard of seven bacteria as our input community. We prepared a sequencing library on the VolTRAX V2 sequence using a MinION mk1b. We basecalled using the latest version of Guppy with the super-accuracy model. We found that increasing read depth benefited each of the assemblers, and nearly complete community member chromosomes were assembled with as little as 10x read depth. Polishing assemblies using Medaka had a predictable improvement in quality. Some assemblers struggled with particular members of the bacterial community, but we found Flye to be the most robust across taxa. We found Flye was the most effective assembler for recovering plasmids. Based on Flye’s consistency for chromosomes and increased effectiveness at assembling plasmids, we would recommend using Flye in future metagenomic studies.

## Introduction

Current methods for sequencing microbes involve isolating and sequencing individual community members, amplicon sequencing 16S rRNA genes, or metagenomics (Garmendía et al. 2012; Petersen et al. 2019). Isolating individual microbes requires culturing, which is often difficult or practically impossible (Garmendía et al. 2012). Sequencing 16S rRNA genes cannot provide information on the entire genomes, such as genes that might increase virulence or provide antibiotic resistance (Petersen et al. 2019). Metagenomics is a method where an entire sample is sequenced, and the individual community members are sorted out later with bioinformatic analyses (Bai et al. 2022). Metagenomic sequencing can detect unculturable and novel community members (Garmendía et al. 2012). The individual community member sequences can be studied to identify pathogens in difficult to diagnose disease, genes that may increase virulence, and look for correlations between co-infecting pathogens that increase disease severity (Garmendía et al. 2012; Kumar et al. 2018; Petersen et al. 2019; Qin et al. 2018). Currently, most metagenomic approaches use Illumina based technology, which produces high accuracy, short reads (Petersen et al. 2019). Over the past several years, Oxford Nanopore Technologies (ONT) has increased sequencing throughput and yield to be reasonable for metagenomic studies. While these reads are error prone, the reads are also orders of magnitude longer than short read platforms (Petersen et al. 2019).

The short reads (150-300 bp) from Illumina sequencing make genome assembly difficult for complex communities. Short read lengths do not facilitate merging multiple contigs built for a genome into a single contig, resulting in fragmented assemblies (Goldstein et al. 2018). Short reads cannot span long repeat regions, causing repeat regions to shrink, providing less complete assemblies (Sevim et al. 2019). More complete genomes can be assembled using long read sequencing technologies, such as ONT or PacBio (Goldstein et al. 2018). ONT sequencers platforms (e.g. MinION) have produced reads greater than 2 mb long and can easily produce libraries with mean read lengths greater than 16 kb, which makes it possible to assemble long repeat regions (Amarasinghe et al. 2020; Payne et al. 2019; Jain et al. 2018). However, the high error rates of nanopore sequencing also prevent short read assemblers from producing quality assemblies with long read data (Jain et al. 2018; Latorre-Pérez et al. 2020).

Three commonly used long read specific assemblers include Flye, Raven, and Redbean (Yang et al. 2021; Latorre-Pérez et al. 2020; Breckell and Silander 2021; Chen, Erickson, and Meng 2020b). Flye is a long read metagenomic assembler that constructs a repeat graph to assemble and polish contigs (Kolmogorov et al.2020, 2019). These contigs are then used to build an assembly graph with A-Bruijn (Kolmogorov et al. 2020, 2019). Previous studies found that while Flye can build more accurate metagenomic assemblies than Raven or Redbean, it also takes more time and memory (Wick and Holt 2019; Latorre-Pérez et al. 2020). Raven is a fast assembler that uses an Overlap-Layout-Consensus (OLC) approach to build an assembly graph from raw reads (Vaser and ikić 2021). For some individual assemblies Raven can have comparable accuracy to Flye after the assemblies are polished, but has less accuracy for metagenomic assemblies (Wick and Holt 2019; Breckell and Silander 2021; Vaser and ikić 2021). Redbean is another fast assembler that follows the OLC concept by using a fuzzy de Bruijn graph to build assemblies from raw reads (Ruan and Li 2020; Rizzi et al. 2019). Previous studies have found that Redbean uses more memory and builds less accurate assemblies than Raven (Wick and Holt 2019; Latorre-Pérez et al. 2020).

Benchmarking is used to compare bioinformatics tools and to determine which tool is best suited for a particular task (Yang et al. 2021; Aniba, Poch, and Thompson 2010). Benchmarking studies for metagenomic assemblers often include well characterized communities or mock communities, like one of the many Zymo-BIOMICS Microbial Community Standards (Latorre-Pérez et al. 2020; Sereika et al. 2021; Goldstein et al.2018; Kolmogorov et al. 2020). Mock communities are synthetic communities composed of multiple known microbes, with known sequences and abundances (Bokulich et al. 2016). This information allows for accurate assessment and comparison of assemblers for metagenomic data

A past benchmarking study using a ZymoBIOMICS Microbial Community Standard found that Raven and Redbean could not build complete assemblies for the *E. coli* and *Salmonella enterica* community members (Latorre-Pérez et al. 2020). Raven did well for the other community members in the ZymoBIOMICS Microbial Community Standard (Latorre-Pérez et al. 2020) Raven also performs well for individual assemblies of *E. coli* (Breckell and Silander 2021; Chen, Erickson, and Meng 2020b). These differences in performance suggest that the high read error rate may cause Raven to confuse genome fragments from other community members with *E. coli* fragments. If so, a higher read accuracy, as produced by the current versions of Guppy, may allow Raven to assemble all community members from the mock community with similar accuracy to Flye. Another weakness of Raven and Redbean, is that they often fail to build assemblies for plasmids (Wick and Holt 2019). These weaknesses may limit the performance of Raven and Redbean for complex metagenomic assemblies, where plasmids may be common and particular community members may be present.

Improvements in converting the electrical signal from nanopore sequencing to to nucleotides (basecalling) have led to increased read level nanopore sequence accuracy. The release of the super-accuracy model to Guppy has pushed modal accuracy to 98% https://nanoporetech.com/accuracy. As the individual reads improve in quality, faster assemblers, such as Raven, may be able to build assemblies of problematic community members, such as *E. coli*, with comparable accuracy to slower, but more accurate assemblers, like Flye.

Here, we compare the completeness and accuracy of metagenomic assemblies built with Flye, Raven, and Redbean. We used data basecalled with the super-accuracy model of Guppy. From this comparison, we contrast the areas of strength and weakness of long read metagenomic assemblers.

## Methods

### Sequencing

We sequenced a mock community standard (ZymoBIOMICS HMW DNA Standard, catalog #D6322) using long read sequencing to compare metagenomic assembly methods. The HMW DNA standard is a synthetic microbial community comprising three gram-negative bacteria, four gram-positive bacteria, and one yeast (Table 1). Bacterial community members have a genome size between 2.73 mb to 6.792 mb, a GC content between 32.9% to 66.2% (Table 1). Each bacterial community also contributed 14% of nucleotides in the mock community (Table 1). The template DNA in the community has a mean length of 24 kb. Sequences can be found at https://s3.amazonaws.com/zymo-files/BioPool/D6322.refseq.zip.

**Table 1:** List of references and community members in the HMW Zymo mock community. No. refers to number.

| NRRL<br>Accession | Organism | plasmids | %<br>GC | Genome size<br>(mb) | Gram<br>stain | %<br>nucleotides | No. genomes |
| --- | --- | --- | --- | --- | --- | --- | --- |
| B-354 | *Bacillus subtilis* | 0 | 43.9 | 04.045 | + | 14 | 13.20 |
| B-537 | *Enterococcus faecalis* | 0 | 37.5 | 02.845 | + | 14 | 18.80 |
| B-1109 | *Escherichia coli* | 1 | 46.7 | 04.875 | - | 14 | 10.90 |
| B-33116 | *Listeria monocytogenes* | 0 | 38.0 | 02.992 | + | 14 | 17.80 |
| B-3509 | *Pseudomonas aeruginosa* | 0 | 66.2 | 06.792 | - | 14 | 07.80 |
| B-4212 | *Salmonella enterica* | 0 | 52.2 | 04.760 | - | 14 | 11.20 |
| B-41012 | *Staphylococcus aureus* | 3 | 32.9 | 02.730 | + | 14 | 19.60 |
| Y-567 | *Saccharomyces cerevisiae* | NA | 38.3 | 12.100 | NA | 02 | 00.63 |

We used 1 ug of the HMW DNA standard as input for the VolTRAX V2 (ONT) to prepare a sequencing library (VSK-VSK002 workflow). The VolTRAX library is analogous to the Rapid Sequencing library and results in additional DNA template fragmentation as the library is prepared. We sequenced the prepared library using the MinION mk1b (ONT) on a r9.4.1 flow cell (FLO-MIN106) for 48 hours (VSK002 script). We basecalled the reads using Guppy version 5.0.7 with the super-accuracy model (-c dna_r9.4.1_450bps_sup.cfg). We set a minimum quality filter of ≥ 10 (-min_qscore 10).

To generate a subsample of reads, we used trycycler (Wick et al. 2021). We used a genome size of 42 mb and the –min read depth parameter to generate subsamples of 420 mb, 840 mb, 1260 mb, 2100 mb, 4200 mb, and 8400 mb. These total yields should theoretically represent 10x, 20x, 30x, 50x, 100x, and 200x read depths. At each read depth, we produced 12 sub-samples for a total of 72 datasets. The mean number of bases, mean longest read length, and mean N50 for each read depth was found using NanoStat –fastq (De Coster et al.2018).

### Assembly and Polishing

For this comparison, we used three commonly used assemblers to construct metagenomic assemblies of our data sets, Flye, Redbean, and Raven. We used Flye version v2.8.3 (Kolmogorov et al. 2020) with default parameters specifying nanopore reads (-nano-raw) and the following options in recover plasmids (-plasmids) and metagenomes (-meta). We used Raven v1.5.1 (Vaser and ikić 2021) with default parameters. We used Redbean v2.5 (Ruan and Li 2020) with default parameters specifying nanopore reads (-x ont), and a genome size of 42 mbases (-g 42m).

We polished all assemblies using one round of Racon v1.4.22 (Vaser et al. 2017) followed by one round of Medaka v1.4.3 https://github.com/nanoporetech/medaka, specifying the super-accuracy model (-m r941_min_sup_g507). For Racon we used the ONT suggested parameters: score for matching bases (-m 8), score for mismatching bases (-x -6), gap penalty (-g -8), window size (-w 500), and mean quality threshold for each window (-q -1).

### Quality assessment

We measured assembly quality and completeness with the genome fraction output by MetaQuast v5.1.0 (Mikheenko, Saveliev, and Gurevich 2016). For MetaQuast we used the references in (Table 1) to measure the completeness of both the polished and unpolished metagenomic assemblies.

We measured assembly accuracy with the median Q-score output by Pomoxis assess_assembly https://github.com/nanoporetech/pomoxis/. Pomoxis was used with the references in (Table 1) to find the quality scores (Q-scores) of the assemblies. For each assembly, we calculated Q-scores for chromosomes and plasmids separately.

We completed all analysis, including assembly, polishing, and assembly quality assessment on a server with an Intel Core i9 9900K 3.6GHz Eight Core (16 thread) CPU, a Nvidia Quadro GV100 gpu, and 128 GB of ram. We measured the time required and the max memory used to build each assembly using GNU time with parameter -f %ee. The time, assembly, polishing, MetaQuast, and Pomoxis steps were automated using custom bash scripts https://github.com/jeremyButtler/assembler-scripts.

We used R v4.1.1.1 (R Core Team, n.d.) with ggplot2 (Wilkinson 2011), cowplot (Wilke 2020), ggpubr (Kassambara 2020), tidyr (Wickham 2021), data.table (Dowle and Srinivasan 2021), stringr (Wickham 2019), and RColorBrewer (Neuwirth 2014) to build graphs for the metagenome fraction, genome fraction, median Q-scores, number of misassemblies, time, and maximum memory usage. The metagenome fraction was found by dividing the number of bases that were aligned to a community member in a replicate by the total bases in the community.

## Results

### Subsampling statistics

We sequenced the ZymoBIOMICS HMW DNA Standard on a nanopore sequencer and subsampled reads into subsamples of 420 mb (~10x read depth), 840 mb (~20x read depth), 1260 mb (~30x read depth), 2100 mb (~50x read depth), 4200 mb (~100x read depth), and 8400 mb (~200x read depth). For each targeted read depth, our mean number of bases was very close to are target number of bases pairs (Table 2). The mean N50 between our read depths only differed by 18 base pairs (15012 to 15030 bp) (Table 2). Each time the read depth was doubled, we saw a two-fold increase in the mean number of reads (Table 2).

**Table 2:** Subsample statistics for each read depth. Each read depth had 12 subsamples. No.bases is the number of bases.

| Read depth | Mean No.reads | Mean N50 | Mean No.base pairs (mb) |
| --- | --- | --- | --- |
| 10x | 45373 | 15012.25 | 419.28 |
| 20x | 91345 | 15020.75 | 838.87 |
| 30x | 137018 | 15020.08 | 1258.72 |
| 50x | 228363 | 15023.42 | 2099.35 |
| 100x | 456726 | 15027.83 | 4198.77 |
| 200x | 913452 | 15030.75 | 8398.78 |

### Chromosome

#### Genome fraction

Across all read depths, we found Flye produced assemblies with near 100% metagenome fractions (Figure 1 (a)). Even at our smallest read depth of 10x, Flye recovered nearly 100% of the metagenomic fraction (Figure 1 (a)). With increasing read depth, Raven and Redbean produced assemblies with improved metagenome fractions (Figure 1 (a)). Raven and Redbean reached a maximum metagenome fraction of 95% at 200x read depth (Figure 1 (b)). At the individual community member level, Raven and Redbean had the most difficulty in the assembly of *Escherichia coli* and *Salmonella enterica*, recovering less than 90% genome fraction even at 200x read depth (Figure 1 (b)).

**Figure 1:**
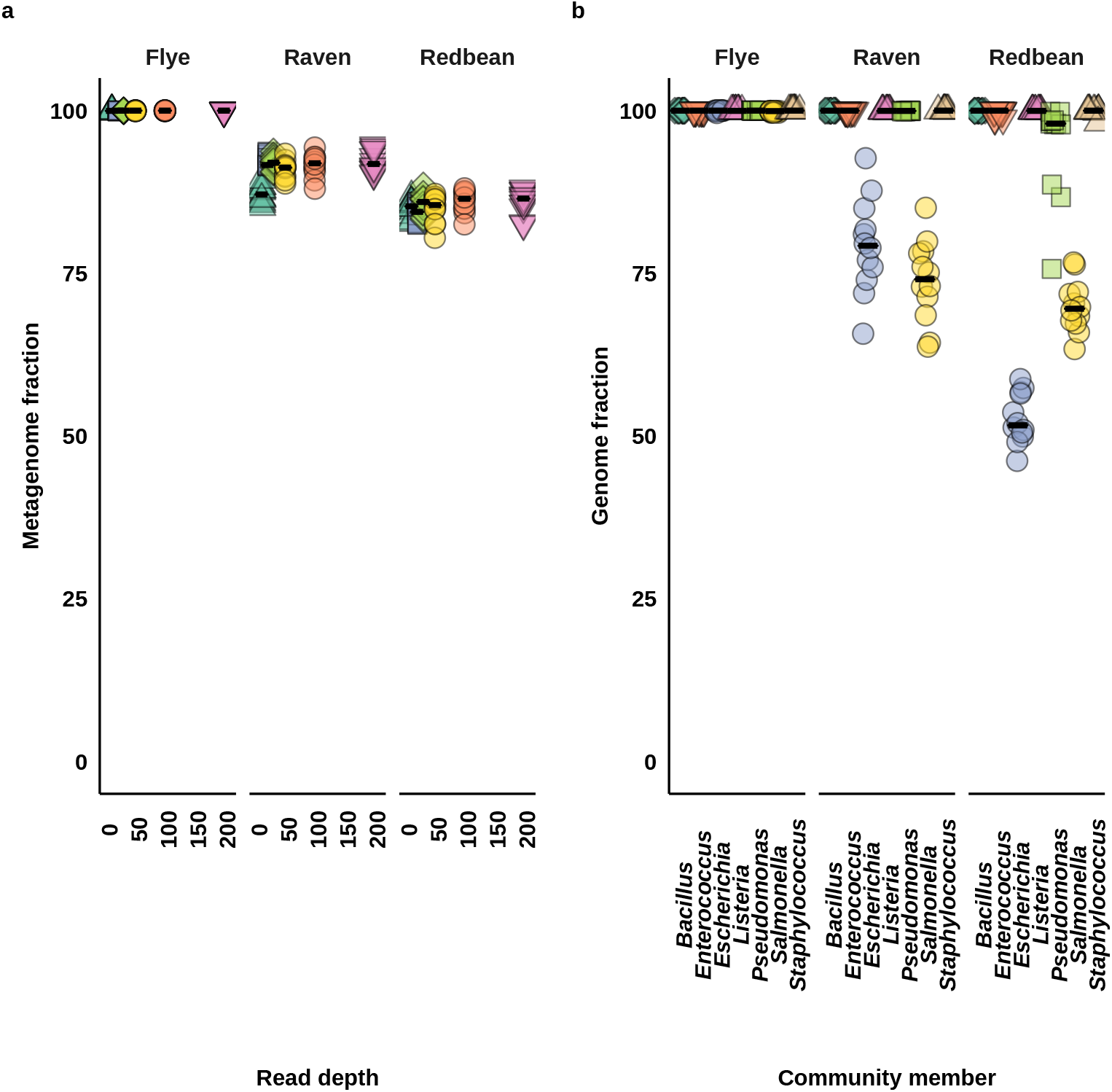
Chromosome completeness. Horizontal bars indicate the medians across replicates. a) The metagenome fractions for all isolates in a replicate. Green circles indicate are polished assemblies. b) The genome fraction for each isolate at 200x read depth. Color and shape indicate different community members.

#### Accuracy (Q-score)

Across all read depths, we found Flye produced the most accurate metagenomic assemblies, followed by Raven, and then Redbean (Figure 2 (a)). Increased read depth and polishing, predictably improved the median quality scores (Q-scores) of assemblies from all assemblers (Figure 2 (a)). All assemblers had a large improvement in Q-scores between 10x and 50x read depth (Figure 2 (a)). At 200x read depth Flye reached a maximum Q-score of 50, while Raven and Redbean reached a maximum Q-score of 46 and 45 respectively (Figure 2 (a)).

**Figure 2:**
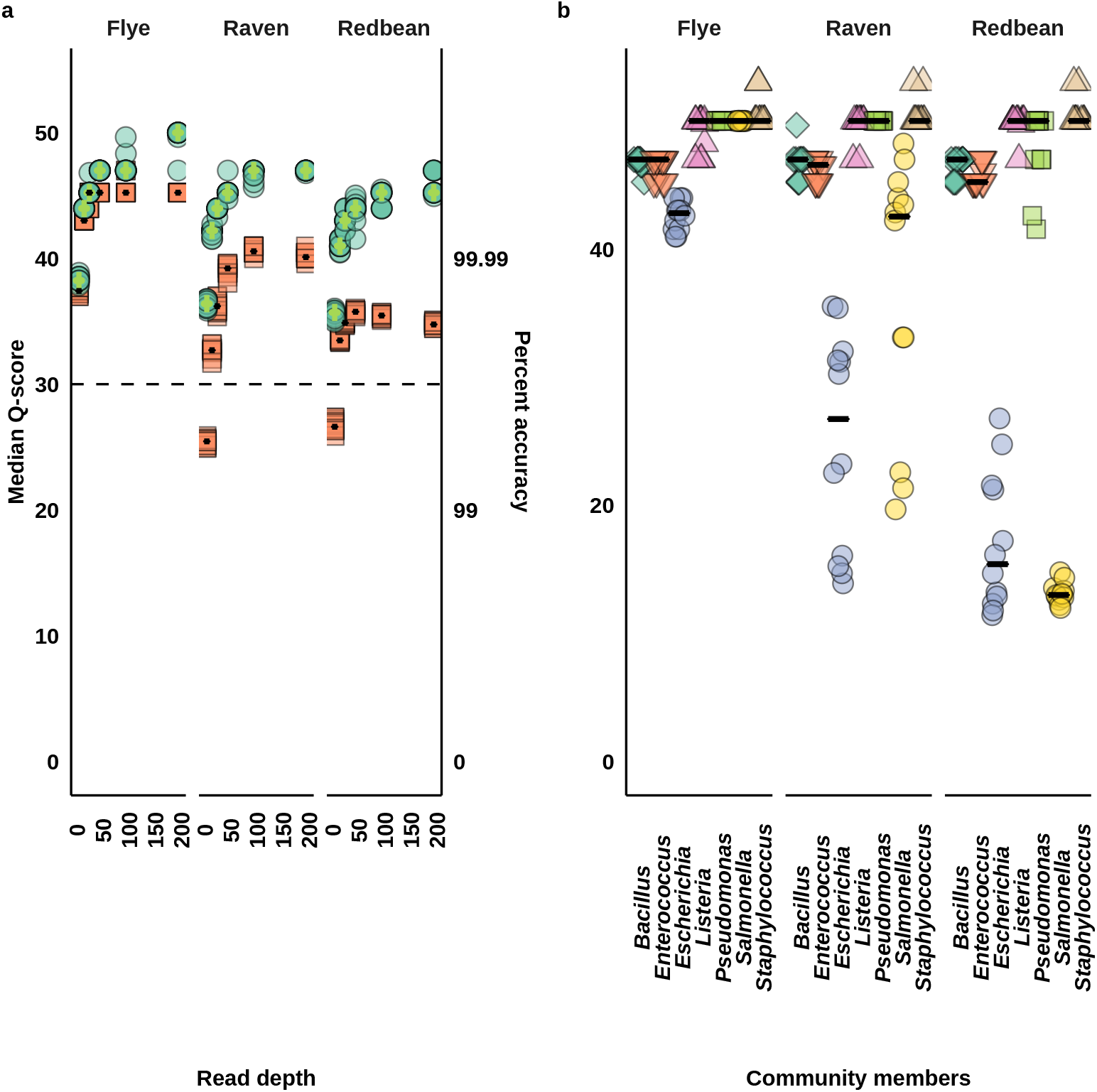
Chromosome accuracy. Horizontal bars indicate the median across replicates. a) Median quality score (Q-score) for chromosomes in each assembly. Green circles and green horizontal bars indicate polished assemblies. b) Median Q-score for each isolate at 200x read depth. Color and shape indicate difference community members.

At the individual community member level, Raven and Redbean had the most difficulty in the assembly of *E. coli* and *S. enterica* (Figure 2 (b)). *E. coli* assemblies produced with Flye were more accurate (median Q-score 42.81) than those from Raven (26.73) and Redbean (under 20) (Figure 2 (b)). *S. enterica* assemblies produced by Flye were highly accurate (median Q-score 50), while Raven was slightly less accurate (42.54), but Redbean produced error prone assemblies (under 20) (Figure 2 (b)). We also found that Raven and Redbean, but not Flye, had over 10 miss-assemblies for *E. coli* and *S. enterica* (Figure S1).

### Plasmids

#### Genome fraction

Across all read depths, we found Flye recovered over 94% of the plasmid genomes (Figure 3 (a)). After 50x read depth Flye recovered nearly 100% of the plasmid genomes (Figure 3 (a)). After 20x read depth, Raven and Redbean decreased the recovery of plasmid genomes (Figure 3 (a)). Raven and Redbean assembled a maximum of 95% of the plasmid genome at 20x read depth (Figure 3 (a)).

**Figure 3:**
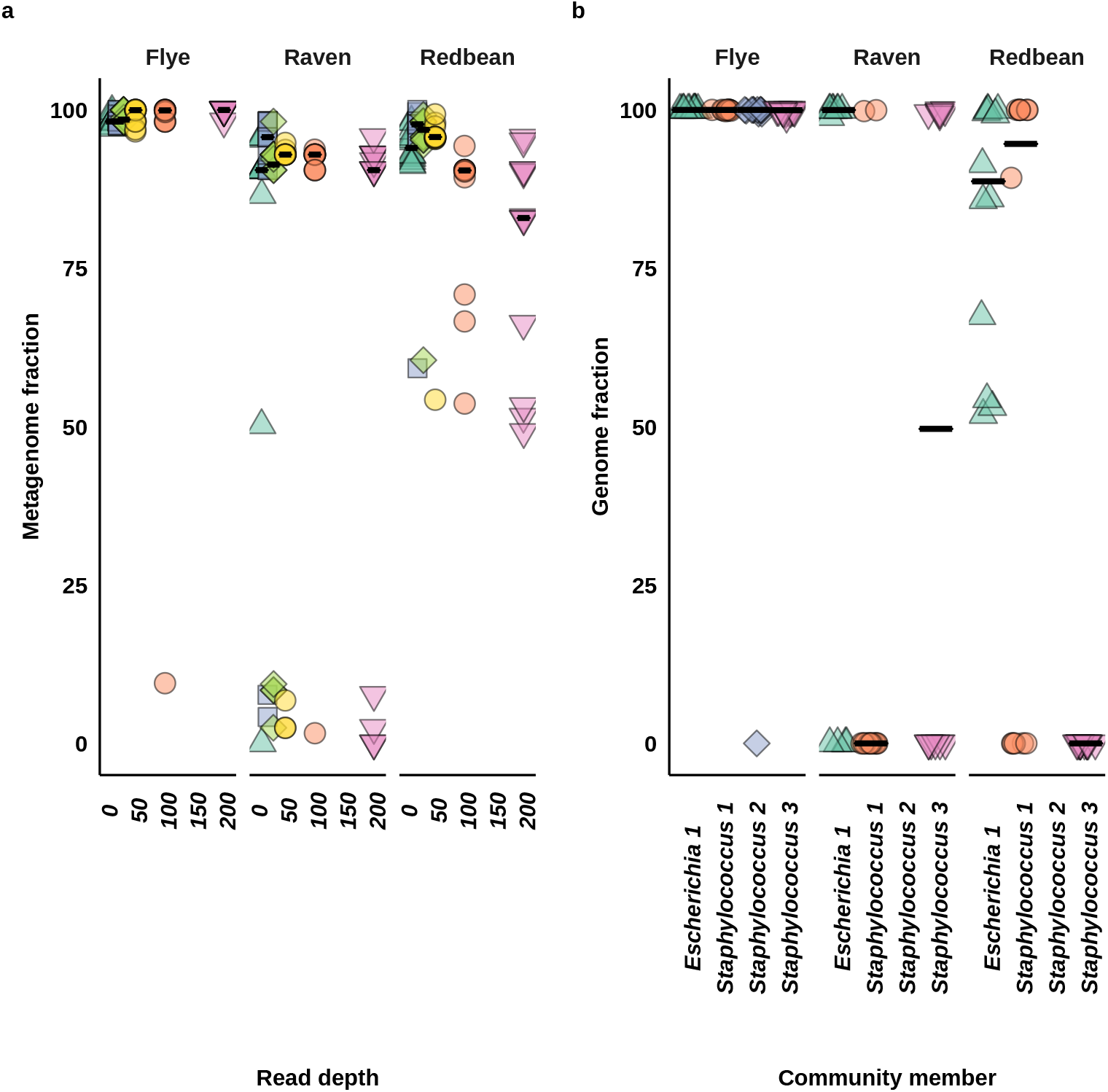
Plasmid completeness. Horizontal bars indicates the median genome fraction across replicates. a) Metagenome fraction for plasmids from each replicate. b) Genome fraction for each plasmid at 200x read depth. *E. coli* is 110009 bases, *S. aureus* 1 is 6339 bases, *S. aureus* 2 is 2218 bases, and *S. aureus* 3 is 2995 bases long.

At the individual plasmid level, Raven and Redbean both struggled with the plasmids smaller than 7 kb (Figure 3 (b)). Raven and Redbean assembled more of plasmids under 7 kb at 30x and 50x read depth than at 200x read depth (Figure S2b; Figure S2a). Raven could assemble the 2995 bp plasmid for all replicates at 50x read depth, but not at 200x read depth (Figure S2b).

#### Accuracy (Q-score)

Across all read depths, we found that found that Flye assembled the most accurate plasmids (Figure 4 (a)). With increased read depth, Flye produced more accurate plasmid assemblies (Figure 4 (a)). However, Polishing did not improve the accuracy of Flye plasmid assemblies (Figure 4 (a)). At 100x read depth, Flye plasmid assemblies had a median Q-score of 50 (Figure 4 (a)).

**Figure 4:**
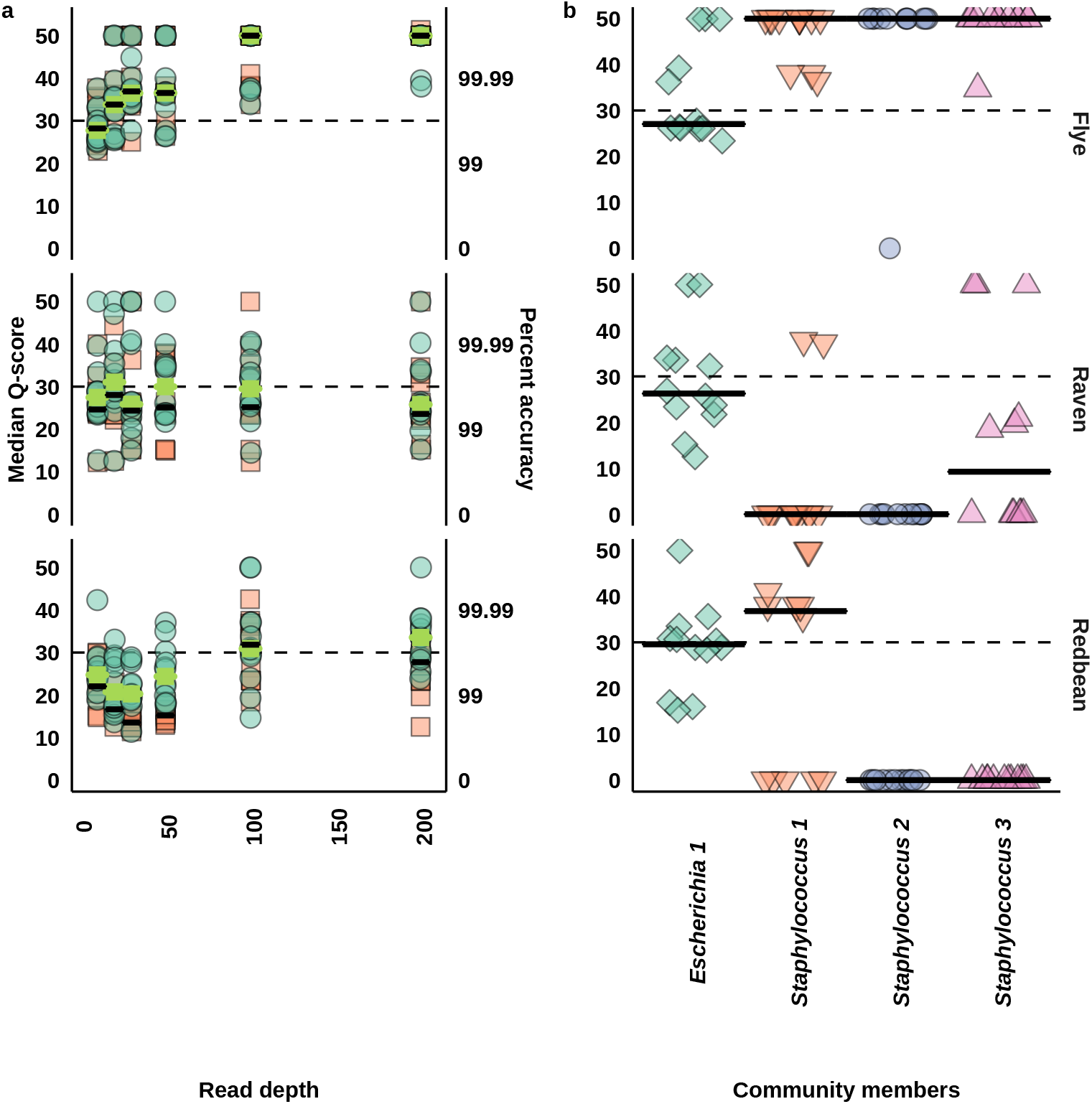
Plasmid accuracy. Horizontal bars indicate the median across replicates. A Q-score of 50 indicates a Q-score of infinity (100% accuracy). a) Median Q-score of all replicates at 10x, 20x, 30x, 50x, 100x, and 200x read depth. Green circles and horizontal green bars indicated polished assemblies. The dashed line indicates the highest Q-score for Raven. b) Median Q-score for each plasmid at 200x read depth. *E. coli* is 110009 bases, *S. aureus* 1 is 6339 bases, *S. aureus* 2 is 2218 bases, and *S. aureus* 3 is 2995 bases long.

Across all read depths, polishing Raven and Redbean plasmid assemblies resulted in more accurate plasmid genomes (Figure 4 (a)). Increased read depth did not imporeve the accuracy of Raven produced plasmid assemblies (Figure 4 (a)). Beyond 50x read depth, Redbean produced more accurate plasmid assemblies than Raven (Figure 4 (a)). However, Raven build more accurate plasmids assemblies than Redbean when the read depth was under 100x (Figure 4 (a)).

At the individual plasmid level, only the *E. coli* 110009 bp plasmid could be assembled by all assemblers (Figure 4 (b)). All assemblers had a similar accuracy for the *E. coli* plasmid (Q-scores around 26) (Figure 4 (b)). All assemblers were able to assemble the *E. coli* plasmid without misassemblies, but Flye and Redbean did have misassemblies for the plasmids under 7 kb (Figure S3). However, Flye assembled almost all replicates for each plasmid and had near perfect median Q-scores for plasmids under 7 kb (Figure 4 (b)).

### Assembly time and memory usage

Predictably, we found that assemblers needed more time and memory to build an assembly with greater read input (Figure 5a and Figure 5b). When the read depth was under 50x, all assemblers used less than 30 minutes to complete an assembly (Figure 5a). At 200x read depth, Flye needed over 400 minutes to complete an assembly (Figure 5a). With that same input Raven required just 50 minutes and Redbean required only 25 minutes to complete an assembly (Figure 5a).

**Figure 5:**
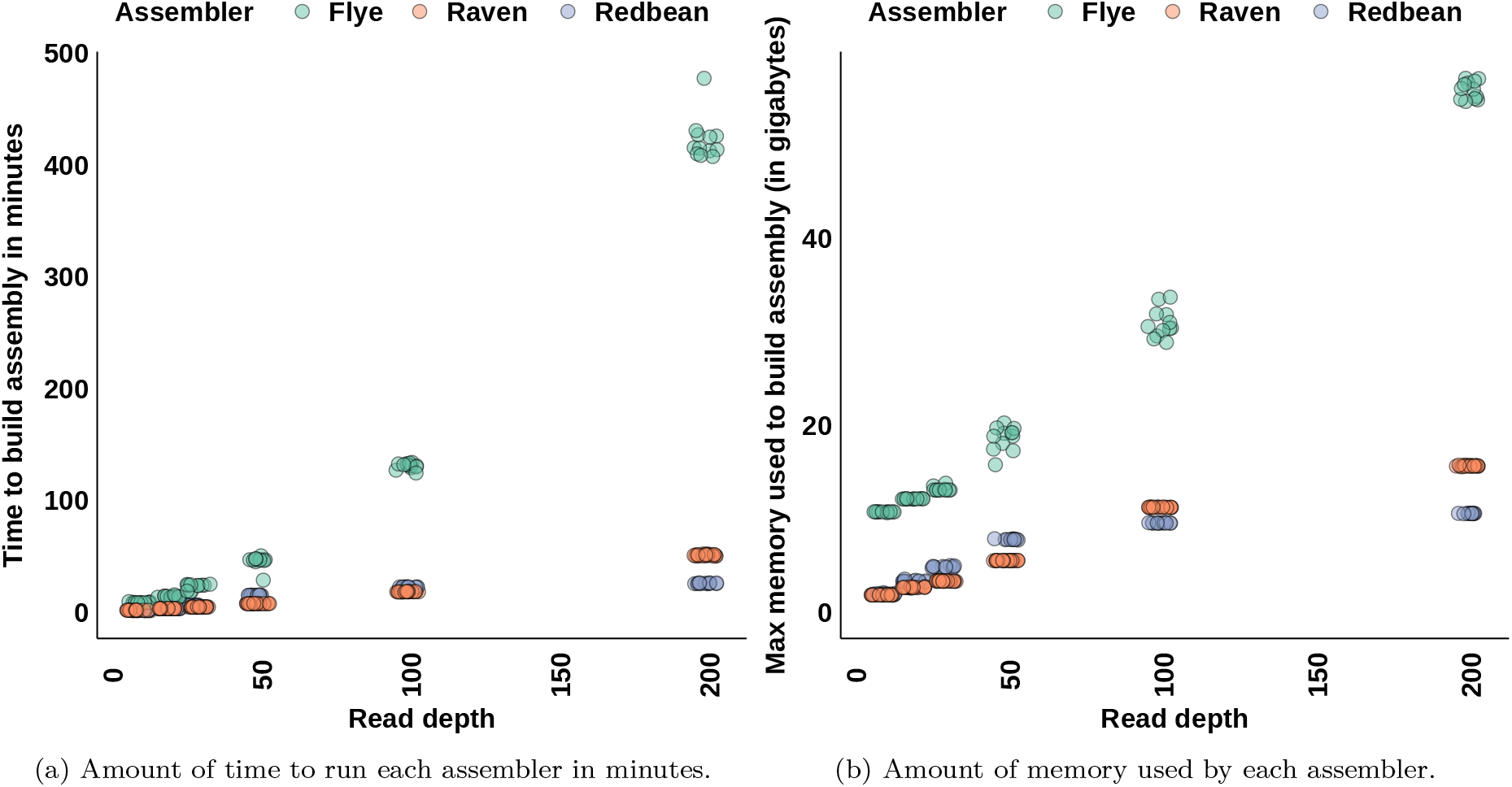
Time and memory usage of each assembler.

Across all read depths, Raven and Redbean used less memory than Flye to build an assembly (Figure 5b). At read depths under 100x, Raven used less memory than Redbean to build an assembly (Figure 5b). At 50x read depth, Raven needed 5.5 Gb of memory to build an assembly, while Redbean needed 7.7 Gb of memory to build an assembly (Figure 5b). Beyond 50x read depth, Raven used more memory than Redbean to build an assembly (Figure 5b). At 200x read depth, Raven used 15.6 Gb of memory to build an assembly, while Redbean used 10.5 Gb of memory to build an assembly (Figure 5b). Flye used the most memory to build an assembly, requiring 10.6 Gb of memory at 10x read depth and 55.8 Gb of memory at 200x read depth (Figure 5b).

## Discussion

We compared the accuracy and completeness of metagenomic assemblies built by three long read assemblers, Flye, Raven, and Redbean. For chromosomes, we found Flye was the only assembler that made near complete and accurate genomes for all community members. For plasmids, we found Flye was the only assembler that could assemble all plasmids reliably. However, Raven and Redbean were superior to Flye in time and memory usage.

### Effect of read depth

For chromosomes, we found with increased read depth, all assemblers made more accurate and complete assemblies. We found that there was a sharp increase in accuracy between 10x and 50x read depth. At 10x read depth, Flye was the only assembler that had near complete metagenome fractions. Showing that Flye should be used for low read depth datasets. However, for more accurate assemblies, future metagenomic studies should continue to aim for a read depth of at least 30x.

For plasmids, we found most plasmids under 7 kb were assembled best by Flye, with the most plasmids recovered at 200x read depth. However, Raven and Redbean had decreased small plasmid recovery at deeper read depths and performed best at read depths between 20x or 50x. The decrease in assembled plasmids under 7 kb at deeper read depths suggests that Raven and Redbean are discarding smaller reads and contigs at deeper read depths. This results in plasmids under 7 kb being missed at deeper read depths but being retained at more shallow read depths. These observations are consistent with Wick and Holt (2019), who also found that both Raven and Redbean struggled to complete assemblies of smaller plasmids. These results highlight the weakness in Raven and Redbean for recovering plasmids.

We found that the accuracy of the larger *E. coli* plasmid (Q-score under 30) was much lower than the chromosome assemblies (40 or 50). This suggests that the plasmids have more error prone regions, assemblers are more likely to make misassemblies for plasmids, or that the plasmid references have more errors than the chromosome references. For reference errors, Flye could often assemble plasmids under 3 kb with no indels or mismatches and with only 2 to 3 misassemblies. Errors in the references are a less likely but still a potential explanation for why Flye, Raven, and Redbean had poor performance for the *E. coli* plasmid.

For misassembly errors, we found all assemblers had no misassembly in *E. coli* plasmid assemblies at 200x read depth, showing that the problem is not from misassemblies in the *E. coli* plasmid. Other sources of errors in the *E. coli* palsmid could be from ore error prone regions in the *E. coli* plasmid or errors inserted by the assemblers in the assembly of the *E. coli* plasmid. During the process of generating these results a new version of Flye was released (v2.9), which included improvements for recovering plasmids and accounts for the improved accuracy of the super-accuracy model. However, more testing with a broader range of plasmid sizes is needed to determine if the errors are from error prone regions or from the assembler.

### Metagenomics and viruses

Though our study examined a mock microbial community mostly consisting of bacterial genomes, our results still provide insights into how reliable each assembler may be for viral metagenomic assemblies. The *E. coli* plasmid in our study is 110 kb long, which is close to or under the size of a large virus, such as the 170 to 190 kb African swine fever virus (ASFV) (Gaudreault et al. 2020). We have previously used Flye to recover ASFV successfully from a metagenomic sample (Kovalenko et al. 2019). While the smaller plasmids in our study are near the size of small viruses, such as porcine circovirus type 2, which is 1.76 Kb long (Breitbart et al. 2017).

For the larger plasmids and likely larger viruses, we found that Raven or Redbean would likely work as well as Flye. However, only Flye could make reliable assemblies for the smaller plasmids and so, is the only reliable assembler for smaller viruses. Even then Flye will often have a few misassemblies, so it might be best to use an assembler, like viralFlye that is designed for viruses (Antipov et al. 2022). However, viralFlye is specialized for virus detection and thus has limitations on the max genome size (Antipov et al. 2022). This may limit viralFlye’s use for bacterial community members. Making Flye or assemblies made with both viralFlye and Flye the best option for sequencing mixed communities of viruses and bacteria.

### Effect of polishing

We found that polishing improved the accuracy of all chromosome assemblies. However, for Flye and Redbean, polishing continued to improve the accuracy at 200x read depth. This suggests that even more data will improve the accuracy of polished Flye assemblies. To achieve highly accurate assemblies, we would recommend polishing and using the greatest read depth as possible.

For Flye, polishing had little effect on the accuracy of plasmid assemblies. Instead, most plasmids smaller than 3 Kb had no indels or mismatches at 200x read depth. This shows that polishing did not decrease the accuracy of the perfect assemblies. Likely, the high accuracy was due to the genome sizes of the plasmids being smaller than the error rate of consensuses assemblies (one error in 10000 bases for chromosomes). The idea of size is somewhat supported by the ten fold larger *E. coli* plasmid assemblies built by Flye having much higher error rates (median Q-score ~ 28) than the plasmids under 3 Kb. Since polishing provides large improvements for chromosomes, while having no decrease in accuracy for plasmids, we would recommend polishing all metagenomic assemblies.

### Problem Isolates

We found that Raven and Redbean struggled to build assemblies of *E. coli* and *Salmonella enterica*. Latorre-Pérez et al. (2020), also found that Raven and Redbean struggled with *E. coli* and *S. enterica* strains for the log and even mock communities from ZymoBIOMICS, both of which use the same *E. coli* and *S. enterica* strains as the HMW DNA Standard Mock Community. However, in a non-metagenomic study, Chen, Erickson, and Meng (2020a) found that Raven could assemble complete genomes for a different strain of *E. coli* and possibly a different serovar of *Salmonella* (*S*. Typhimurium). This suggests that either the strain of *E. coli* used in the mock community is a problematic strain or that assembling genomes of *E. coli* combined wit *S. enterica* is difficult. Breckell and Silander (2021) found that strain specific characteristics of different *E. coli* made some *E. coli* strains harder to for assemblers to assemble, so it is possible that the strain of *E. coli* in the mock community could be a more difficult strain to assemble. However, Breckell and Silander (2021) found that problematic strains of *E. coli* were problematic for all assemblers. Flye had few misassemblies for *E. coli* at 200x read depth and had more accurate assemblies of *E. coli* than Raven or Redbean. This evidence is not consistent with a problematic strain of *E. coli*. However, we cannot fully eliminate the idea that the strain of *E. coli* in the mock community may be a more difficult strain to assemble.

### Other studies

To the best of our knowledge, our study is the first study to compare metagenomic assemblies made by Flye, Raven, and Redbean using supper-accurate basecalled reads. We found Flye still made more accurate and complete genomes than Raven or Redbean when used with highly accurate reads. This is consistent with a previous comparison of Flye, Raven, and Redbean assemblies made from the less accurate reads (Latorre-Pérez et al. 2020). Like Sereika et al. (2021) we found accurate genomes could be built from read depths as low as 30x using Flye (Q 45 at 30x). This is an improvement from the Q-score of 43.6 at 80x read depth seen by Broddrick et al. (2020). We also know from Sereika et al. (2021) that even higher accuracies can be achieved if a r 10.4 flow cell is used instead of a r 9.4 flow cell.

Like Breckell and Silander (2021) and Latorre-Pérez et al. (2020), we found Flye and Raven to be better than Redbean in assembling complete genomes. However, unlike Breckell and Silander (2021), but like Latorre-Pérez et al. (2020), we found Flye assembled more accurate assemblies than Raven. The difference may be that Breckell and Silander (2021) looked at assembling single isolates instead of metagenomes like us and Latorre-Pérez et al. (2020). This suggests that Raven may be better suited for assembling single isolates than metagenomics.

Like Wick and Holt (2019), we found Flye needed more time and memory than Raven and Redbean to complete an assembly. The large time and memory demands of Flye may limit Flye to lab use or at least limit Flye to high end laptops. However, Flye was the only assembler able to assemble the entire mock community at Q-scores greater than 40. Also, the use of the super-accuracy basecalling model will likely require a higher end laptop with a good GPU. This makes the high time and memory usage of Flye less of an issue.

### Summary

We found Flye was more reliable than Raven or Redbean for building accurate and complete assemblies for both chromosomes and plasmids from metagenomic communities. We found that Raven and Redbean struggle to recover small plasmids. This suggests that Flye would be a better choice for assembling viral community members. For our study’s community, Raven and Redbean only performed better than Flye in the computational resources needed to build an assembly. However, for a metagenomic study using the super-accurate basecalling model, the extra time and memory usage needed to run Flye would likely be minimal. On the other hand, the cost in accuracy from problematic communities members or missing small plasmid and virus assemblies from Raven and Redbean could lead to misinterpretations. Thus, for future metagenomic studies that use the super-accurate basecalling model, we would recommend using Flye.

## Acknowledgements

We are thankful to Tracie Haan and Taylor Seitz who sequenced the synthetic community used in our study. We would like to thank members of Drown Lab, Olin Silander, Ursel Schütte, Diane Wagner, Karsten Hueffer, and Eric Bortz who provided feedback on the experimental analysis and manuscript drafts. Research reported in this publication was supported the Department of Biology and Wildlife, by Alaska INBRE, an Institutional Development Award (IDeA) from the National Institute of General Medical Sciences of the National Institutes of Health under grant number P20GM103395 as well as Alaska BLaST which is supported by the NIH Common Fund, through the Office of Strategic Coordination, Office of the NIH Director with the linked awards: TL4GM118992, RL5GM118990, UL1GM118991.

## Data Availability

The sequencing data for this project can be found in the NCBI SRA https://www.ncbi.nlm.nih.gov/sra/ under accession number PRJNA903965.

## Supplemental figures

**Figure S1:**
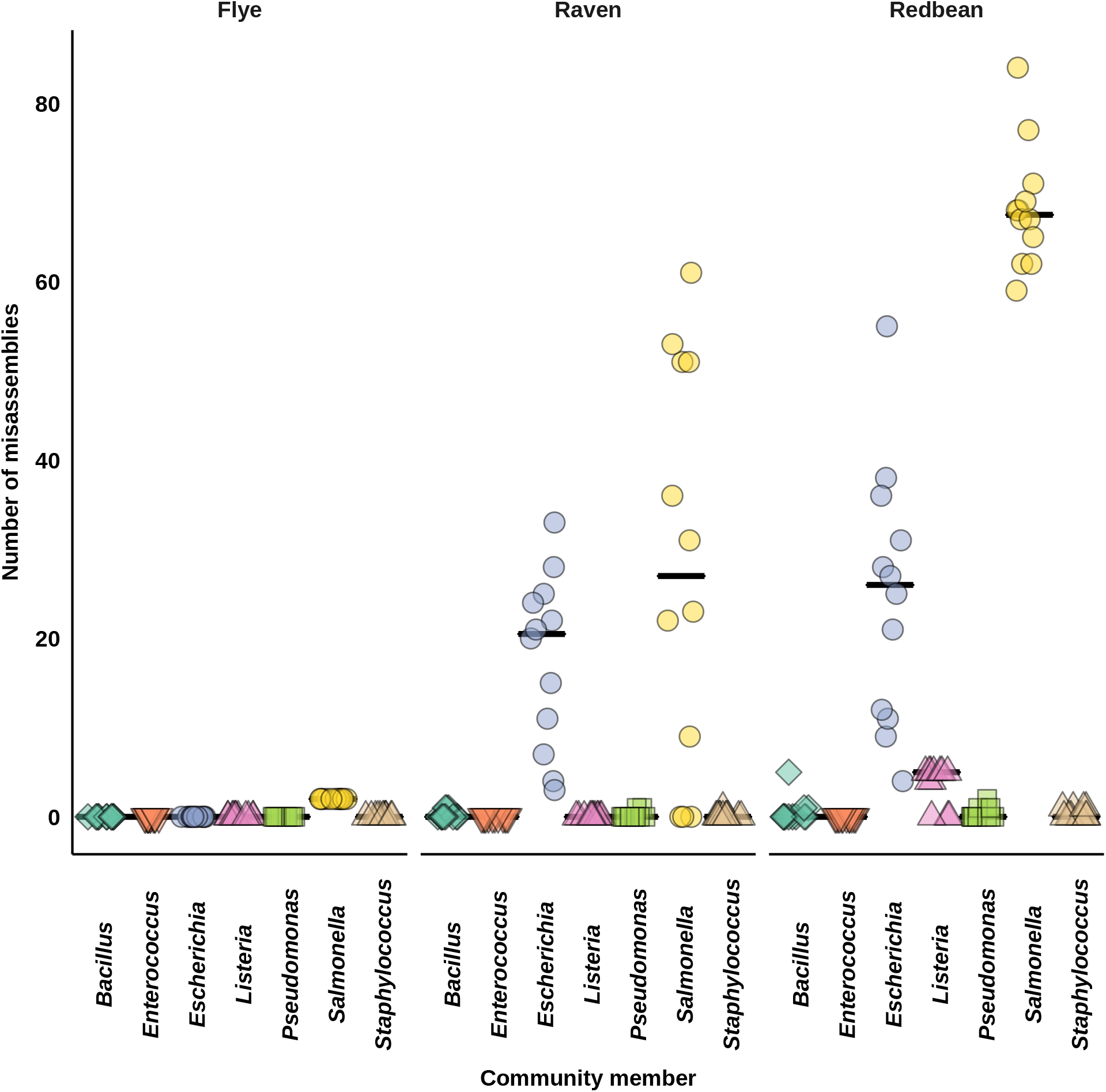
Chromosome misassemblies at 200x read depth. Horizontal bars indicate the median value across replicate samples.

**Figure S2:**
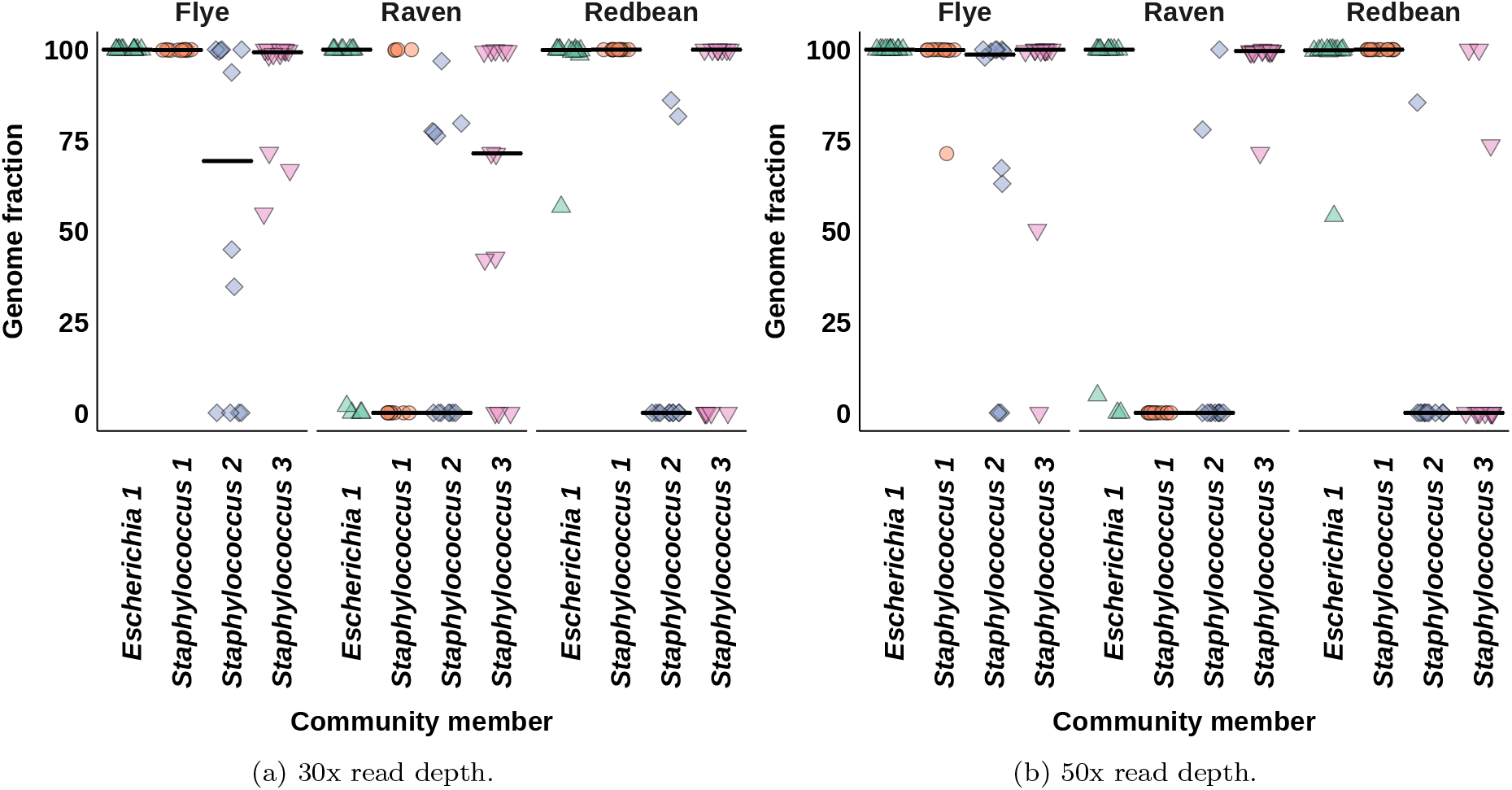
30x and 50x plasmid completeness. Horizontal bars indicate the median across replicates. *E. coli* is 110009 bases, *S. aureus* 1 is 6339 bases, *S. aureus* 2 is 2218 bases, and *S. aureus* 3 is 2995 bases long.

**Figure S3:**
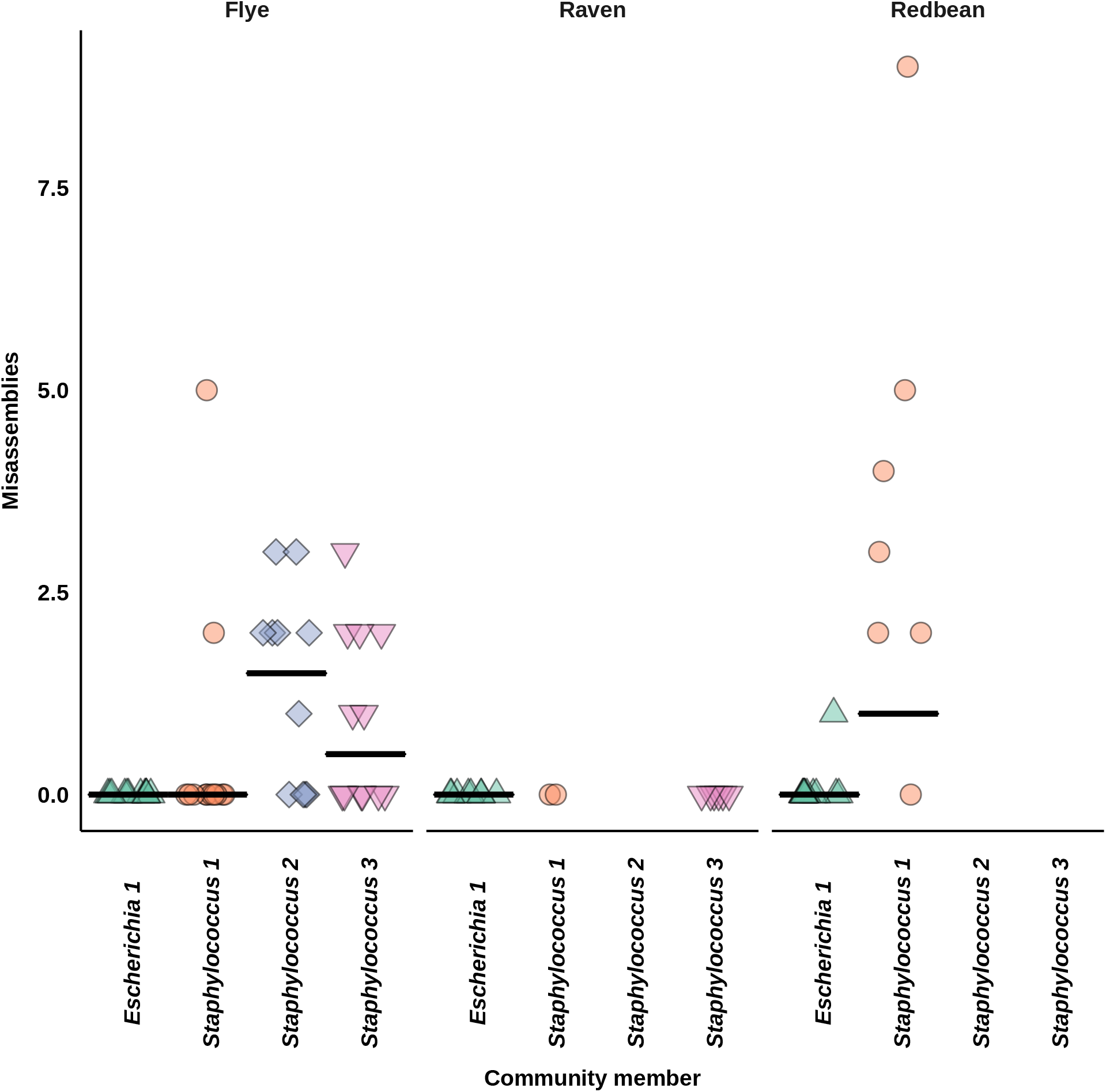
Plasmid misassemblies at 200x read depth. Horizontal bars indicate the median value across replicate samples.

